# FADU: A Feature Counting Tool for Prokaryotic RNA-Seq Analysis

**DOI:** 10.1101/337600

**Authors:** Matthew Chung, Ricky S. Adkins, Amol C. Shetty, Lisa Sadzewicz, Luke J. Tallon, Claire M. Fraser, David A. Rasko, Anup Mahurkar, Julie C. Dunning Hotopp

## Abstract

**Motivation:** The major algorithms for quantifying transcriptomics data for differential gene expression analysis were designed for analyzing data from human or human-like genomes, specifically those with single gene transcripts and distinct transcriptional boundaries that extend beyond the coding sequence (CDS) as identified through expressed sequence tags (ESTs) or EST-like sequence data. Some eukaryotic genomes and all, or nearly all, bacterial genomes require alternate methods of quantification since they lack annotation of transcriptional boundaries with EST or EST-like data, have overlapping transcriptional boundaries, and/or have polycistronic transcripts.

**Results:** An algorithm was developed and tested that better quantifies transcriptomics data for differential gene expression analysis in organisms with overlapping transcriptional units and polycistronic transcripts. Using data from standard libraries originating from *Escherichia coli* and *Ehrlichia chaffeensis,* and strand-specific libraries from the *Wolbachia* endosymbiont wBm, FADU can derive counts for genes that are missed by HTSeq and featurecounts. Using the default parameters with the *E. coli* data, FADU can detect transcription of 51 more genes than HTSeq in union mode and 21 genes more than featurecounts, with 42 and 18 of these features being ≤300 bp, respectively. Due to its ability to derive counts for otherwise unrepresented genes without overstating their abundance, we believe FADU to be an improved tool for quantifying transcripts in prokaryotic systems for RNA-Seq analyses.

**Availability and implementation:** FADU is available at https://github.com/adkinsrs/FADU. FADU was implemented using Python3 and requires the PySAM module (version 0.12.0.1 or later).

## 1. Introduction

A typical analysis pipeline for a gene expression analysis of transcriptomics sequencing data involves: (a) mapping sequencing reads to a whole genome transcriptome assembly with an aligner like Bowtie (Langmead, et al., 2009), BWA (Li and Durbin, 2009), HISAT (Kim, et al., 2015), or STAR (Dobin, et al., 2013); (b) counting reads or fragments for each gene with a tool like HTSeq (Anders, et al., 2015) or Subread featurecounts (Liao, et al., 2014); and (c) finding differentially expressed genes through the use of tools like DESeq (Anders and Huber, 2010) and edgeR (Robinson, et al., 2010). Most of these tools were designed to analyze human data, and as such, they carefully consider important issues that affect these analyses, such as transcript splicing. However, important and relevant genomic features in other organisms complicate transcriptomics analyses in ways unaddressed with this human-centric focus, for example the polycistronic transcripts of bacterial operons.

Most commonly identified in prokaryotes, operons are transcriptional units that encode polycistronic transcripts with multiple coding sequences (CDSs). This allows for the coordinated transcription and regulation of all the genes in an operon. As an example, the *lac* operon encodes a permease for transporting lactose into the cell and a β-galactosidase which converts lactose to galactose and glucose, allowing for the cis-regulation of multiple functional related genes under a single promoter (Lewis, 2013).

Presently, the two most popular tools for transcriptome analyses are HTSeq (Anders, et al., 2015) and Subread featurecounts (Liao, et al., 2014). Although in most cases, both tools have no issue quantifying transcripts for specific genes, issues arise when a single fragment can be assigned to multiple genes. By default, both HTSeq and featurecounts bin these reads as ambiguous, rather than assigning them to a specific gene. While this may not be as significant of a problem in eukaryotic systems, the features of a prokaryotic genome, namely the smaller gene size, smaller genome size, and the presence of operons, make it difficult for HTSeq and featurecounts to quantify smaller genes, especially those within operons that are smaller than the library insert size.

Here, we test how operons and polycistronic transcripts confound HTSeq and featurecounts, leading to a lack of sequencing data for small genes within operons. We developed a new tool, Feature Aggregate Depth Utility (FADU) to quantify transcription in bacterial genomes. We test FADU on multiple bacterial genomes to demonstrate its utility at capturing sequence data for these underrepresented genes.

## 2. System and Methods

### 2.1 Availability of data sets

Three data sets were used in all analyses consisting of RNA-Seq paired-end data from standard, non-stranded libraries originating from (a) *E. coli* and (b) *E. chaffeensis* and stranded libraries from (b) wBm. The sequencing reads for the three datasets can be found in the NCBI Sequencing Read Archive at the following accession numbers, respectively: (pending), SRX485438, and SRX2505171.

### 2.2 FADU, featureCounts, and HTSeq comparisons

For comparative analyses, FADU was run using –count_by fragment and all other default options. HTSeq vO.lO.O (Anders, et al., 2015) was run using default settings while changing the mode for mode-specific analyses. Subread featureCounts v1.6.1 (Liao, et al., 2014) was run using the -p option to specify counting by fragments and/or-0 or-fractional to specify counting different methods of counting ambiguous reads depending on the analysis. Unrooted dendrograms were generated using the R package APE v5.0 (Analysis of Phylogenetics and Evolution) (Paradis, et al., 2004; Popescu, et al., 2012). Bootstrap values were obtained using the R package pvclust v2.0-0 (Suzuki and Shimodaira, 2006). The principal component analysis was performed using the R packages FactoMineR v1.39 (Le, et al., 2008) and factoextra vl.0.5 (http://sthda.com/english/rpkgs/factoextra/).

## 3. Algorithm

### 3.1 Creating a mapping index using an annotation file

A mapping index is created that contains each position in the reference genome. For each feature present in the GFF3 or GTF annotation input file, coordinates are marked in the mapping index for each of the features’ positions. If the reads are ‘stranded’ or ‘reverse-stranded’, a separate mapping index is created and marked for each strand. Each of these coordinates are marked using the features’ attribute id. At positions shared by multiple features, the position will be marked as an overlap between two features. These positions, along with positions absent of any feature, will be excluded from downstream feature count calculations. From this, a statistics file will be written that contains the following information for each feature: (a) strand, (b) length of feature, (c) number of coordinates mapping solely to that gene, (d) proportion of non-overlapping coordinates compared to length of feature.

### 3.2 Calculating read/fragment counts for each feature

For each BAM file, the read depth is calculated using the depth function of samtools with the -aa option If FADU is set to calculate fragment depth, all non-properly-paired reads are discarded by default and the read depth is adjusted to determine the fragment depth at all positions. The user can elect to keep all mapped read (as opposed to properly paired reads), including singletons and discordant reads, in which case all reads will be included in the fragment depth totals. To calculate the fragment depth from the samtools depth output, for each of the properly-paired reads, all coordinates in the insert region between the paired reads are incremented by one and coordinates where the reads overlap are decremented by one. If BAM data is identified as “stranded” or “reverse-stranded”, each BAM file is split into a “(+)-stranded” and a “(-)-stranded” BAM file, based on the bitwise flag field in the input BAM file. Each stranded BAM will have its read or fragment depth calculated separately.

For each input BAM file, the average read length or average fragment length is determined to calculate counts for each feature. If the option to keep only properly paired reads is set, then only those reads will factor into the average read or fragment length calculations.

For each feature, all the coordinates that mapped solely to this feature are collected. The total depth of this feature is calculated by summing the read or fragment depth for each coordinate collected in the feature, and this total is divided by the average read or fragment length to derive a fragment count for each feature. The feature ID and count statistic is written to a file. If multiple BAM files were used as input, then the counts of each individual input will be written to a separate file.

## 4. Implementation

FADU was written entirely using the Python3 programming language. It relies heavily on the PySam module (version 0.12.0.1 or later) to parse information from the BAM alignment files, to write intermediate BAM files, and to perform basic samtools commands. The program supports multiprocessing, and the user can specify the number of processes to be utilized. Each process will handle a separate BAM input file if a list of files is provided. FADU was tested in the UNIX environment.

To minimize the amount of memory used, temporary files are written when possible to keep track of read depth and the coordinates of properly paired reads. In addition, when read depth is converted into fragment depth, only the bases with nonzero depth are read into memory.

## 5. Results

### 5.1 Gene detection performance of FADU, HTSeq, and featureCounts

To assess how FADU compares to featureCounts and HTSeq in deriving counts, we used paired-end sequencing data from three different sets of transcriptome data: (a) paired end reads from a standard (i.e. not strand-specific) library constructed from *Escherichia coli* RNA, (b) paired end reads from a standard library constructed from *Ehrlichia chaffeensis* RNA, and (c) paired-end reads from a strand-specific library constructed from *Wolbachia* endosymbiont of *Brugia malayi* wBm RNA.

Of the 4,647 protein-coding genes detected in *E. coli,* counts for 51 genes could be obtained using FADU, but not HTSeq union, the default HTSeq mode **(Figure 2a).** Because HTSeq union discards fragments spanning multiple features, in the case when unstranded data is being used, HTSeq union is likely unable to identify these genes because: (1) the gene is largely overlapping another feature either on the same or opposite strand or (2) the gene is within an operon and smaller than the average library fragment size. Because FADU calculates counts based on the depth at only positions unique to any given feature, FADU can assign partial counts to multiple features per fragment, allowing for the increased representation of smaller genes, as well as the unique portion, if any, of overlapping genes. Supporting this, 42 of the 51 genes unable to be detected with HTSeq union are ≤300 bp in size **(Table 1).** While HTSeq union and featureCounts are largely similar, featureCounts handles ambiguous reads differently. Given a fragment that maps to multiple features, featureCounts will assign the paired-end fragment to the feature that maps to the majority of the individual paired-end reads (Liao, et al., 2014). When comparing FADU with featureCounts. 21 genes were only detected using FADU, with 18 of these genes being ≤300 bp in size **(Figure 2b).**

**Table 1.** Key Properties of Data Examined

| **Species** | ***E. coli*** | ***E. chaffeensis*** | **Bm** |
| --- | --- | --- | --- |
| Strand-Specificity | no | no | reverse |
| Number of Sequenced Paired-End Reads | 215,149,159 | 46,817,709 | 75,945,674 |
| Number of Mapped Paired-End Reads | 184,454,369 (85.7%) | 3,132,709 (6.7%) | 351,928 (0.5%) |
| Genes | 4,647 | 1,002 | 1,006 |
| Genes Detected in FADU but not HTSeq or featureCounts | 51 | 5 | 31 |

Similarly, FADU can derive counts for an additional five genes in *E. chaffeensis* compared to HTSeq union or an additional two genes when compared to featureCounts. All genes detected only with FADU in *E.chaffeensis* were ≤300 bp in length. With wBm, 31 additional genes were detected in FADU when compared to HTSeq union, of which 10 are ≤300 bp, while 24 additional genes were detected when compared to featurecounts, of which **7** are ≤300 bp **(Figure 2ab).** This indicates that despite featurecounts being able to detect a greater number of genes than HTSeq union, FADU can derive counts for genes that neither HTSeq or featurecounts can by default.

HTSeq has two additional modules to derive counts for transcriptome data that both attempt to assign ambiguous reads. In the case that a fragment overlaps multiple features, HTSeq intersection-nonempty takes the intersect of the features found at each non-empty position and if only one feature is returned, a count is assigned to that feature. Similarly, HTSeq intersection-strict takes the intersect of the features found at all positions of the fragment and again, if only one feature is returned, a count is assigned to that feature (Anders, et al., 2015). While this allows for the assignment of more ambiguous fragments, smaller genes are still under-represented. Additionally, because HTSeq intersection-strict also discards fragments that partially map to intergenic regions, and because most prokaryotic organisms currently have no UTR annotations, this will result in discarding reads at the 5’-and 3’-end of prokaryotic transcripts. In all cases, for genes smaller than the library insert size, it becomes difficult to extract any meaningful fragment counts.

When comparing FADU to HTSeq intersection-strict, FADU derives counts for an additional 182 genes. HTSeq intersection-strict fails to obtain counts for >100 additional genes compared to HTSeq union **(Supplementary Figure 1ab),** confirming the inability of HTSeq-intersection-strict to accurate assess prokaryotic transcriptome data for instances in which the reference has limited UTR annotations. Supporting this, HTSeq intersection-strict fails to detect an additional 60 genes in *E. chaffeensis* and 71 genes in wBm when compared to FADU. HTSeq intersection-nonempty performs similarly to HTSeq union, failing to detect 48, 4, and 31 genes when compared to FADU in *E. coli, E. chaffeensis,* and wBm, respectively, indicating regardless of which module used, HTSeq is too conservative in assigning reads to genes.

**Figure 1:**
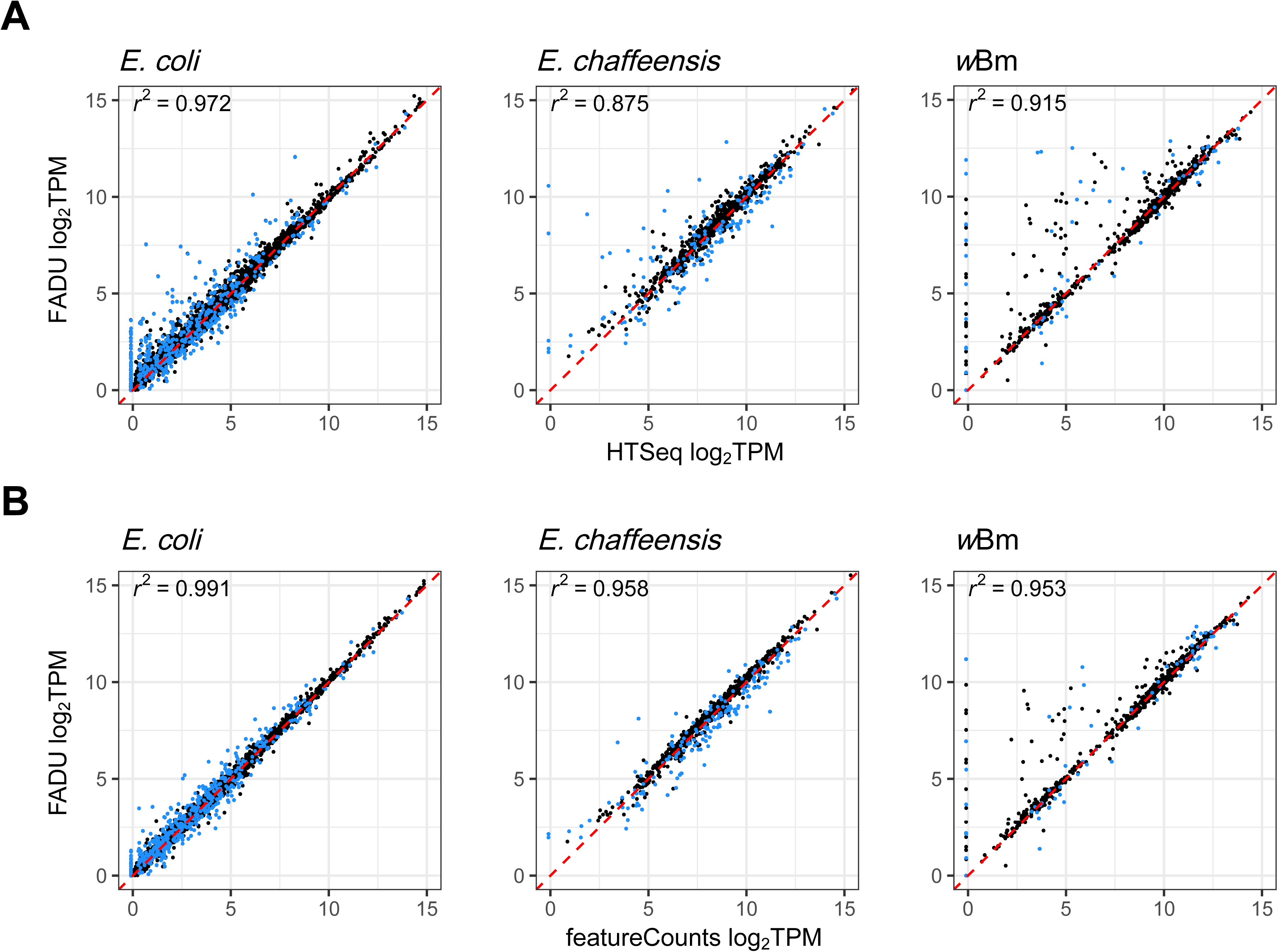
Comparison of TPM values derived from FADU to HTSeq union and featureCounts default For three different sets of RNA-Seq paired-end data from *E. coli, E. chaffeensis,* and wBm, the log_2_ TPM values for genes quantitated using FADU were plotted against the log_2_ TPM values for genes quantitated with **(A)** HTSeq union and **(B)** featureCounts default. Each point is representative of a single gene, with points in blue being representative of genes <300 bp in length. Genes with similar count values are expected to lie close to the identity line (x=y; red). Genes whose expression values are more elevated in FADU lie above the identity line while genes whose expression values are elevated in HTSeq of featureCounts lie below the identity line. Genes able to be quantified in FADU but not in HTSeq union or featureCounts default lie on the y-axis. These genes include very highly transcribed genes suggesting that they are missed by all the tools except FADU, and not that they are poorly transcribed, small genes.

While featurecounts does not have any distinct modules, there are two options which help to assign counts for ambiguous reads. The first is the -O option, in which cases where a fragment overlaps multiple features, a single count is added to both. The second is specifying both the -O and the - fragment options, in which case fragments that overlap *x* features are given a count of *1/x.* For the *E. chaffeensis,* and wBm datasets, FADU obtains counts for the same number of genes as both featurecounts overlap and featurecounts fractional-overlap **(Supplementary Figure 2ab).** However, in the *E. coli* dataset, both modes of featureCounts have counts for nine additional genes compared to FADU. Of these nine *E. coli* genes, eight are completely overlapped by another gene either on the same or opposite strand. Because these genes have no unique positions with which FADU can use to determine count values, FADU returns a fragment count of 0 for these genes. The last gene, E2348C_0713, is 642 bp long with the first 104 bp being overlapped by another gene. At most, featureCounts overlap gives E2348C_0713 a fragment count of 2, while featureCounts fractional-overlap assigns a count of 1, indicating that only two fragments map to E2348C_0713 map within the first 104 bp. Because FADU calculates fragment counts using only unique positions of a gene, FADU assigns a fragment count of 0 to E2348C_0713.

**Figure 2:**
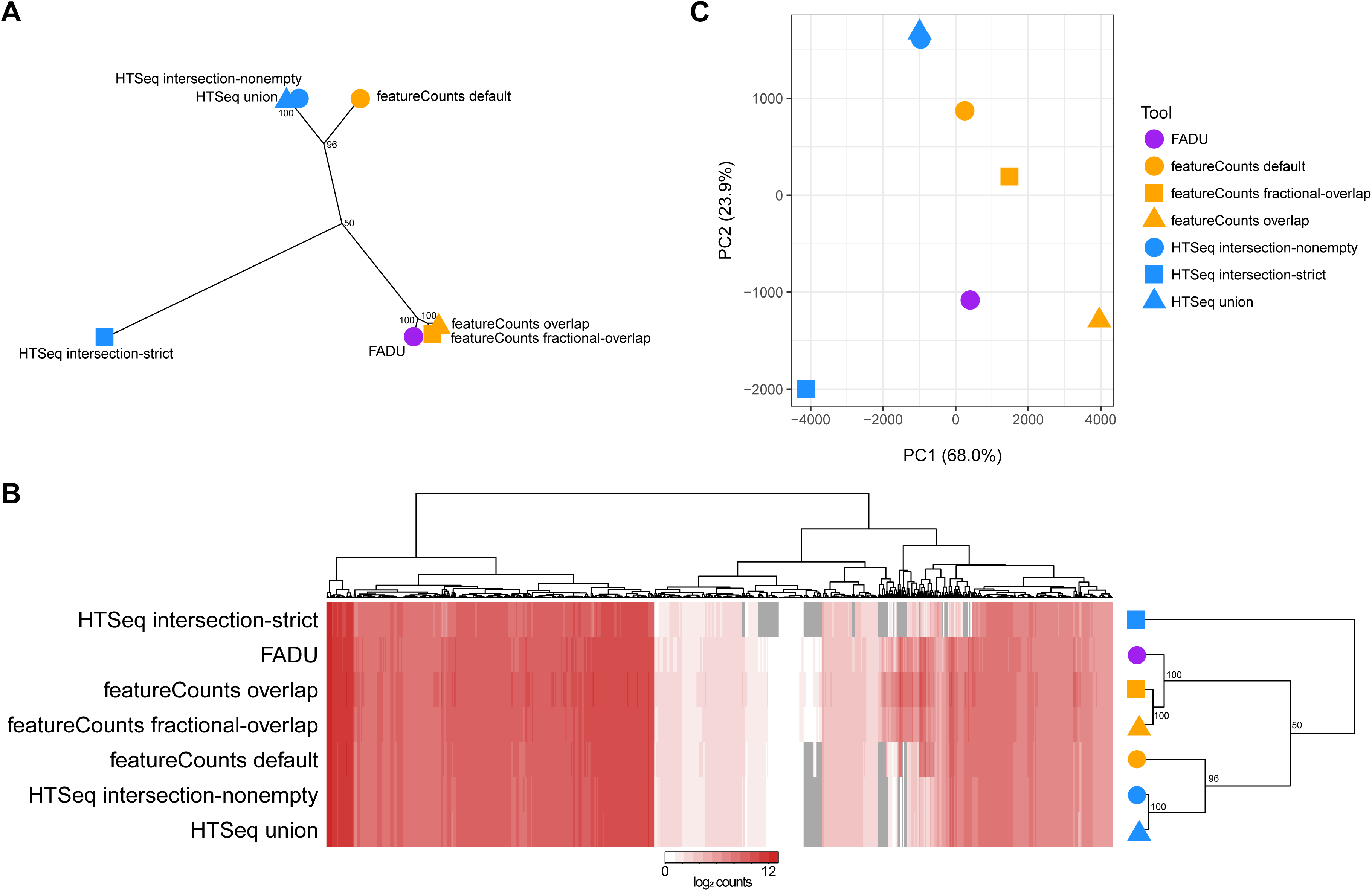
Clustering patterns of the different count values in ivBm derived with HTSeq modules, featureCounts modes, and FADU **(A)** An unrooted dendrogram with 1000 bootstraps was generated using the log_2_ count values from wBm calculated using HTSeq, featureCounts, and FADU. The dendrogram reveals three distinct clusters of (1) featureCounts default, HTseq union, and HTSeq intersection-nonempty; (2) HTSeq intersection-strict; and (3) FADU, featureCounts overlap, and featureCounts fractional-overlap. **(B)** The log_2_ count values for all wBm genes with count values derived from at least one of the tools was used to generate a heatmap. The wBm genes are displayed on the horizontal axis while each of the tools are displayed on the vertical axis. All cells in grey describe genes with no count value in its corresponding tool. Bootstrap values for both the unrooted and squared dendrograms are located next to their corresponding nodes. **(C)** A principal component analysis for all wBm count values derived from each of the tools was done. Each color corresponds to either FADU, HTSeq, or featureCounts, while each shape represents the specific mode of the tool used.

### 5.2 Comparative analysis of FADU, HTSeq, and countFeatures in ***w*Bm**

Using the wBm dataset, we sought to determine the similarity of FADU compared to each of the different modes of HTSeq and featureCounts. Fragment count values from the three HTSeq modules, three featureCounts modes, and FADU were used for a clustering analysis. An unrooted dendrogram of the different tools shows three distinct groups that cluster on how each of the different tools handle fragments mapping to multiple features **(Figure 3a).** FADU, featureCounts overlap, and featureCounts fractional-overlap, which are more liberal in assigning counts, form a cluster while HTSeq union, HTSeq intersection-nonunique, and featurecounts default, which are all more conservative, form another cluster. HTSeq intersection-strict clusters with neither of the groups, due to it being the most stringent in assigning fragment counts to features.

A heatmap showing counts from each of the eight tools shows HTSeq intersection-strict to have the greatest number of genes with no assigned counts **(Figure 3b).** Only genes with derived count values from at least one tool are shown. The cluster containing featurecounts default, HTSeq union, and HTSeq intersection non-empty contain slightly less genes with no assigned count values while the cluster containing FADU, featurecounts overlap and featureCounts fractional-overlap contain the least. Although featureCounts overlap is able to assign count values to the same number of genes as FADU, it over-counts genes by assigning a full count value to all genes overlapped by a single fragment. In the case of a fragment overlapping a two gene operon, featureCounts overlap would assign a full count to both, despite there only being a single fragment. To diminish over-counting, featureCounts fractional-overlap instead assigns a fractional count value based on the number of features a fragment overlaps. While this alleviates the issue, featureCounts fractional-overlap implies that all features overlapped by a fragment contribute equally to the fragment, which may not necessarily be true. The problem is particularly acute if the overlap is a relatively small fraction of the feature. FADU assigns count values based on the percentage of the fragment that is overlapped, such that a higher partial read count is assigned to the gene with the greater overlap. By doing so, FADU can assign higher counts from ambiguous fragments to genes that the fragment most likely originated from, while still being able to derive counts for smaller genes.

A principal component analysis of the counts show less discrete clusters compared to those seen in the unrooted dendrogram **(Figure 3c).** While the counts from HTSeq union and HTSeq intersection-nonempty are grouped together, no other two counts cluster closely with another. In the first principal component, which accounts for 68.0% of the variation observed in the counts, the top 20 contributing genes are primarily represented by genes with lower counts in the three HTSeq modes relative to featureCounts and FADU **(Supplementary Figure 3a).** Similarly, the top 20 contributing genes in the second principal component, which accounts for 23.9% of the variation observed, separates the HTSeq-derived counts from the featureCounts and FADU counts **(Supplementary Figure 3b).** In both principal components, there are genes with lower counts in HTSeq intersection-strict relative to all other counts, reflecting the conservative nature in which it assigns counts. Because of how featureCounts overlap derives counts, it will always have greater than or equal to the highest number of counts relative to all other algorithms tested.

## 6. Discussion

During transcript quantification for RNA-Seq analyses, the handling of fragments that overlap multiple features must be addressed. This may not be as much of an issue in many eukaryotes, where genes are larger and spaced further apart. But in prokaryotes, the closer proximity of genes coupled with the presence of operons leads to a large number of fragments being classified as ambiguous. Tools such as HTSeq and featureCounts have different modules and/or options to handle these ambiguous fragments, but smaller genes, especially those in operons, become either under-or over-represented depending on the tool. In this study, we present FADU, a novel tool for transcript quantification in RNA-Seq analyses that addresses these issues.

While it can be easy to think of all Illumina data as being equal, our analysis suggests that small genes near or below the insert size of the library are specifically being lost. This bears more scrutiny and consideration in prokaryotic transcriptomic sequencing projects, since the insert size of the library varies between samples and is not frequently reported. Our results suggest that these small genes could be differentially reported, in a purely artefactual way, during feature counting and impacts downstream analyses, like differential expression, clustering, and PCA-type analyses.

Importantly, FADU is not a counting algorithm and as such does not report counts as other algorithms have over the past several years. As such, it does not return integers, instead returning fraction-based rational numbers. As such the output of FADU cannot be used in downstream tools that require integer counts, such as some differential expression analysis tools. It can, however, be used with success in edgeR and in calculating TPMs and z-scores. There may, however, be a new need for further development of statistical analysis tools that do not require integer-based data.

Compared to the default HTSeq and featureCounts modes, which largely discard ambiguous reads, FADU assigns partial read counts based on the percentage of the fragment that is within the unique positions of gene. By doing this, FADU is able to assign partial counts to features that are missed by both HTSeq and featureCounts by default. While HTSeq and featureCounts have options that allow for the assignment of reads to these features, we find that both the overlap and fractional-overlap options overstate their abundance, especially in the case of completely overlapped genes. FADU weighs the percentage of each fragment covered by a feature so in the case that a fragment does overlap multiple features, instead of assigning equal counts to both features, partial read counts are assigned based on the percentage of the fragment covered by the feature. Due to its ability to derive counts for otherwise unrepresented genes without overstating their abundance, we believe FADU to be an improved tool for quantifying transcripts in prokaryotic systems for RNA-Seq analyses.

## Acknowledgements

This project was funded in part by federal funds from the National Institute of Allergy and Infectious Diseases, National Institutes of Health, Department of Health and Human Services under grant number U19 AI110820.

---

**Supplementary Figure 1: Comparison of TPM values derived from FADU to HTSeq intersection-nonempty and intersection-strict**

For each of the three different sets of RNA-Seq paired-end data from *E. coli, E. chaffeensis,* and *w*Bm, the log_2_ TPM values for genes quantified using FADU were plotted against the log_2_ TPM values for genes quantified with two of the non-default HTSeq modules: **(a)** HTSeq intersection-nonempty and **(b)** HTSeq intersection-strict. Each point is representative of a single gene, with points in blue being representative of genes ≤300 bp in length. Genes with similar count values are expected to lie close to the identity line (x=y; red). Genes whose expression values are more elevated in FADU lie above the identity line while genes whose expression values are elevated in the HTSeq counterpart lie below the identity line. Genes able to be quantified with FADU but not in HTSeq lie on the y-axis.

**Supplementary Figure 2: Comparison of TPM values derived from FADU to featureCounts overlap and fractional-overlap**

For each of the three different sets of RNA-Seq paired-end data from *E. coli, E. chaffeensis,* and wBm, the log_2_ TPM values for genes quantitated using FADU were plotted against the log_2_ TPM values for genes quantitated with two different featureCounts runs, (a) The first set of plots are run with the featureCounts option overlap, in which multiple genes overlapped by the same fragment are both assigned full counts, **(b)** The second set of plots are run with the featureCounts option overlap and fractional, in which multiple genes overlapped by the same fragment are assigned fractional counts depending on the number of features overlapped by the fragment. Each point is representative of a single gene, with points in blue being representative of genes ≤300 bp in length. Genes with similar count values are expected to lie close to the identity line (x=y; red). Genes whose expression values are more elevated in FADU lie above the identity line while genes whose expression values are elevated in its featureCounts counterpart lie below the identity line. Genes able to be quantified in featureCounts overlap or fractional-overlap but not FADU lie on the x-axis.

**Supplementary Figure 3: Clustering of the twenty top contributing *w*Bm genes in the first and second principal components**

Two heatmaps were generated to visualize the top contributing in **(a)** the first and **(b)** the second principal components analysis of the variation in counts for wBm genes derived using HTSeq, featureCounts, and FADU. For each of the two principal components, the top twenty contributing genes to the variation observed are shown. The horizontal axis of the heatmap describes the tool used while each of the genes are indicated on the vertical axis. The log_2_ count values are shown in each of the corresponding cells.

